# Antigen-dependent IL-12 signaling in CAR T cells promotes regional to systemic disease targeting

**DOI:** 10.1101/2023.01.06.522784

**Authors:** Eric Hee Jun Lee, Cody Cullen, John P. Murad, Diana Gumber, Anthony K. Park, Jason Yang, Lawrence A. Stern, Lauren N. Adkins, Gaurav Dhapola, Brenna Gittins, Wen Chung-Chang, Catalina Martinez, Yanghee Woo, Mihaela Cristea, Lorna Rodriguez-Rodriguez, Jun Ishihara, John K. Lee, Stephen J. Forman, Leo D. Wang, Saul J. Priceman

## Abstract

Chimeric antigen receptor (CAR) T cell therapeutic responses are hampered by limited T cell trafficking, persistence, and durable anti-tumor activity in solid tumor microenvironments. However, these challenges can be largely overcome by relatively unconstrained synthetic engineering strategies, which are being harnessed to improve solid tumor CAR T cell therapies. Here, we describe fully optimized CAR T cells targeting tumor-associated glycoprotein-72 (TAG72) for the treatment of solid tumors, identifying the CD28 transmembrane domain upstream of the 4-1BB co-stimulatory domain as a driver of potent anti-tumor activity and IFNγ secretion. These findings have culminated into a phase 1 trial evaluating safety, feasibility, and bioactivity of TAG72-CAR T cells for the treatment of patients with advanced ovarian cancer (NCT05225363). Preclinically, we found that CAR T cell-mediated IFNγ production facilitated by IL-12 signaling was required for tumor cell killing, which was recapitulated by expressing an optimized membrane-bound IL-12 (mbIL12) molecule on CAR T cells. Critically, mbIL12 cell surface expression and downstream signaling was induced and sustained only following CAR T cell activation. CAR T cells with mbIL12 demonstrated improved antigen-dependent T cell proliferation and potent cytotoxicity in recursive tumor cell killing assays *in vitro* and showed robust *in vivo* anti-tumor efficacy in human xenograft models of ovarian cancer peritoneal metastasis. Further, locoregional administration of TAG72-CAR T cells with antigen-dependent IL-12 signaling promoted durable anti-tumor responses against both regional and systemic disease in mice and was associated with improved systemic T cell persistence. Our study features a clinically-applicable strategy to improve the overall efficacy of locoregionally-delivered CAR T cells engineered with antigen-dependent immune-modulating cytokines in targeting both regional and systemic disease.

## Introduction

Chimeric antigen receptor (CAR)-engineered T cell therapies against solid tumors have largely failed to achieve the robust and durable responses observed in hematologic malignancies^1,2^. Several key distinctions exist between these malignancies, including both physical and immunological aspects, which may be driving the limited responses seen to date in solid tumor CAR T cell clinical trials. These challenges have inspired next-generation strategies to optimize potent and selective CAR molecules, T cell functionalities, routes of T cell administration, and therapeutic combinations to improve solid tumor CAR T cell therapies^3-6^.

Several key improvements to CAR molecules have been developed^4,7,8^. First, the antigen-binding domains, mainly single chain variable fragments (scFvs), have been affinity-tuned to selectively target low or high tumor antigen density^9,10^. These considerations factor in off-tumor antigen expression patterns that may cause unwanted safety issues. The extracellular spacer domain has been shown to regulate T cell persistence through Fc-mediated clearance mechanisms, and its length and conformation can further impact potency and selectivity of CAR T cells^11-13^. While studies have shown that varying transmembrane domains primarily regulate CAR expression stability, more recent evidence supports this domain in regulating CAR signaling^14,15^. It is widely supported that the co-stimulatory domain, typically CD28 or 4-1BB, can greatly impact CAR T cell functionality and *in vivo* expansion and persistence for durable anti-tumor responses^16-18^. Identification of the optimal CAR molecule alone, though, has proven insufficient for producing durable responses in solid tumors and has called for CAR-extrinsic innovation to further augment therapeutic responses. Engineered immune-modulating cytokines or chimeric-switch receptors have been shown to enhance therapeutic activity, although these often unconditional strategies retain the risk of off-target signaling and toxicities and warrant further optimization^3,19-21^.

To address potential off-target toxicities and improve the immediate on-target activity of CAR T cell therapies for solid tumors, local or regional routes of administration compared to systemic intravenous delivery are being explored. Locoregional delivery of T cell therapies is an effective strategy to overcome limitations of cell trafficking, tissue distribution, and on-target off-tumor toxicities by localizing anti-tumor activity at sites of disease. We and others have demonstrated improved anti-tumor responses following locoregional intraventricular delivery of CAR T cells for the treatment of recurrent adult glioblastoma, pediatric brain tumors, and brain metastasis and/or leptomeningeal disease^22-26^. Likewise, intrapleural administration of CAR T cells has been effective in targeting mesothelioma and other pleural diseases^27,28^. Direct local intratumoral administration has been utilized to test CAR T cells targeting tumor antigens with unknown safety profiles^29^. Regional intraperitoneal delivery of CAR T cells, by several groups including our own, has demonstrated improved anti-tumor activity against peritoneal metastasis compared with intravenous delivery^30,31^. However, each of these locoregional delivery approaches is accompanied with a potential risk of limited distribution outside of the regional space and thereby restricts potential targeting of systemic multi-metastatic disease. Several groups have incorporated engineering approaches to improve persistence and trafficking of T cells^3,21,32^, which is likely paramount to targeting widespread metastatic disease following locoregional delivery of cell therapies.

In this study, we focused on expanding our development of tumor-associated glycoprotein-72 (TAG72)-directed CAR T cells for regional targeting of peritoneal cancer metastasis^31^. We empirically optimized our TAG72-CAR construct identifying that the CD28 transmembrane coupled with a 4-1BB co-stimulatory domain greatly improved T cell functionality and yielded durable complete responses *in vivo*. These studies have led to a new phase 1 trial evaluating safety, feasibility, and bioactivity of TAG72-CAR T cells for the treatment of patients with advanced ovarian cancer (NCT05225363). Functional assessment revealed interferon gamma (IFNγ) as a key mediator of CAR T cell cytotoxicity and recursive tumor cell killing *in vitro*, and further validated the role of IL-12 in potent IFNγ induction and downstream CAR T cell activity. Additionally, we engineered a membrane-bound form of IL-12 (mbIL12) with an optimal transmembrane domain that improved potency of CAR T cells *in vitro* and *in vivo*.

Mechanistically, mbIL12 cell surface expression and downstream signaling was antigen-dependent, requiring T cell activation through CAR or T cell receptor (TCR) to promote both *cis* and *trans* signaling. Further, we showed that locoregionally-administered TAG72-CAR T cells with antigen-dependent IL-12 signaling promoted durable control of peritoneal disease and also enhanced peripheral CAR T cell expansion and persistence that eradicated systemic disease. These data not only critically inform on our clinical development program, but also broadly support the use of antigen-dependent immune-modulating cytokines in promoting regional to systemic disease targeting with other CAR T cell therapies and for multiple disease indications.

## Results

### CD28 transmembrane in TAG72-CARs containing a 4-1BB costimulatory domain enhances anti-tumor activity *in vitro*

We recently generated and preclinically evaluated second-generation TAG72-specific CAR T cells containing a 4-1BB intracellular costimulatory domain, which demonstrated potent anti-tumor activity using human xenograft peritoneal ovarian tumor models^31^. The decision to redesign the CAR molecule for optimal functionality was based on our preclinical studies showing a lack of curative anti-tumor activity as well as early phase 1 data using first-generation TAG72-CAR T cells^33^ demonstrating anti-idiotype antibody production in patients, likely contributing to a lack of durable therapeutic responses. First, we re-assessed the antigen-binding single chain variable fragment (scFv) domain of our TAG72-CAR construct in attempts to minimize the potential for anti-CAR immunogenicity and improve T cell persistence. We utilized two additional scFvs (v15, and v59-15: a fusion between v15 and v59) based on the original humanized CC49 scFv (IDEC) that through affinity maturation showed reduced potential for anti-idiotype immunogenicity^34,35^. Two of three scFvs exhibited similar high binding affinities toward TAG72 antigen (IDEC, K_D_ = 33 ± 20 nM; v15, K_D_ = 35 ± 10 nM; v59-15, not determined). For all related *in vitro* and *in vivo* studies, we use human ovarian cancer cell lines that are TAG72-negative (OVCAR8) or are varying in cell surface expression levels of TAG72 (OVCAR3, OV90, and OVCAR8-sTn) (**Supplemental Figure 1**). We incorporated these scFvs into the same CAR backbone we originally published^31^and evaluated their anti-tumor activity *in vitro* and *in vivo* (**Supplementary Figure 2**). TAG72-CAR T cells with the v15 scFv were the most optimal in terms of anti-tumor activity as compared with IDEC and v59-15.

Next, we generated seven v15 scFv-based TAG72-CAR constructs with varying extracellular spacer domains and lengths (termed EQ, dCH2, CD8h, HL, and L), transmembrane domains (CD4tm, CD8tm, CD28tm), and intracellular costimulatory domains (CD28, 4-1BB) (**Figure 1a**). While all seven CAR molecules comparably expressed CD19t marker, we saw higher cell surface CAR expression with Fc-derived spacers (EQ, dCH2), as measured by Protein L staining of the scFv, which closely matched the staining pattern when anti-Fc antibodies to detect the extracellular spacer domain were used (**Figure 1b**). We next evaluated cytotoxicity function of these TAG72-CAR T cells using *in vitro* co-culture killing assays against cancer cell lines with varying TAG72 expression. In general, we found that TAG72-CAR T cells containing the dCH2 spacer domain showed superior functionality with the greatest tumor cell killing, highest CD137 activation and enhanced antigen-dependent T cell proliferation, and robust IFNγ and IL-2 cytokine production (**Figure 1c-e and Supplemental Figure 3**). The three TAG72-CAR leads (dCH2(28tm)28z, dCH2(4tm)BBz, and dCH2(28tm)BBz) showed highest antigen density-dependent T cell activation and cytokine production. Additionally, we showed the greatest PD-1 exhaustive phenotype in CD28 costimulatory domain-containing CAR T cells, in line with previous reports using other CARs^16,36-38^. Unexpectedly, while TAG72-CAR variants with CD8h and HL spacers showed an apparent lack of CAR cell surface expression, they showed varying tumor cell killing potential with some capacity, albeit suboptimal, to induce antigen-dependent T cell proliferation. Finally, we showed the lowest PD-1 expression and highest IFNγ production with our TAG72-CAR T cells that contained the CD28tm and 4-1BB costimulatory domain. We also evaluated short-term signaling pathways following antigen stimulation in the three TAG72-CAR leads. While the 4-1BB costimulatory domain-containing CAR T cells demonstrated reduced downstream PLCy, SLP76, and ERK signaling as compared with CAR T cells containing the CD28 costimulatory domain, the addition of CD28tm to 4-1BB costimulation partially restored downstream signaling (**Figure 1f-g**). These findings suggest that transmembrane domains of CARs have the potential to regulate early downstream signaling following CAR stimulation and can greatly impact CAR T cell anti-tumor functionality.

**Figure 1.**
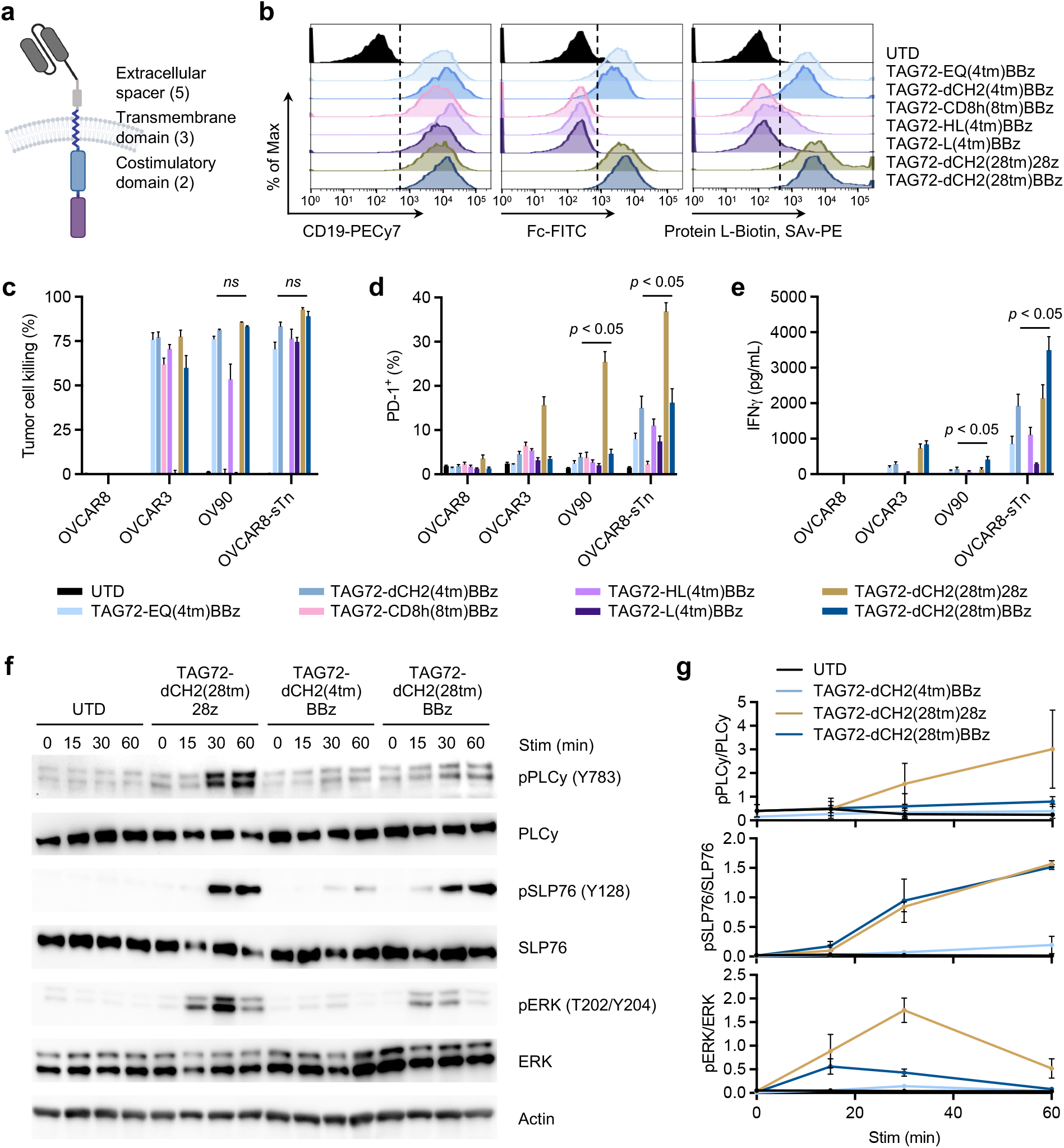
CD28 transmembrane in CARs containing a 4-1BB costimulatory domain enhances anti-tumor activity *in vitro*. **(a)** Diagram of the lentiviral construct with TAG72-CAR containing the humanized scFv targeting TAG72, varying extracellular spacer domains (EQ, dCH2, CD8h, HL, L), transmembrane domains (CD4tm, CD8tm, CD28tm), and intracellular costimulatory domains (4-1BB, CD28) followed by a cytolytic domain (CD3z). A truncated non-signaling CD19 (CD19t), separated from the CAR sequence by a ribosomal skip sequence (T2A), was expressed for identifying lentivirally transduced T cells. **(b)** Untransduced (UTD) and 7 different TAG72-CAR T cells positively enriched for CD19t were evaluated by flow cytometry for CD19t expression to detect lentiviral transduction of CARs (b), fragment constant (Fc) derived spacer containing CARs (c), or Protein L to detect the scFv (d). **(c-e)** *In vitro* tumor cell killing activity relative to UTD (c), expression of PD-1 (d), and IFNγ production by ELISA (e), of CAR T cells against tumor targets (TAG72-OVCAR8; TAG72+ OVCAR3, OV90, and OVCAR8-sTn) after 24 hr (for ELISA) or 72 hr of co-culture at an effector:target (E:T) ratio of 1:4. (**f**) Western blotting analysis of early downstream signaling mediators following CAR T cell stimulation of indicated TAG72-CAR T cells. (**g**) Quantification of band density of phosphorylated proteins levels over their respective total protein levels.

### CD28 transmembrane domain in 4-1BB-based TAG72-CAR T cells induces durable therapeutic responses *in vitro* and *in vivo*

From these studies, we proceeded to further “stress-test” challenge the three TAG72-CAR T cell lead candidates. We first evaluated the cytotoxicity potential of CAR T cells over an extended 10-day co-culture assay with OV90 tumor cells. Using xCELLigence as a readout, TAG72-CAR T cells containing the CD28tm and 4-1BB costimulatory domain displayed the greatest anti-tumor activity (**Figure 2a**). We then performed recursive tumor cell killing assays by rechallenging TAG72-CAR T cells with OV90 tumor cells. We observed two intriguing patterns; both 4-1BB costimulatory domain-containing CARs showed superior antigen-dependent T cell expansion profiles, whereas the CD28 transmembrane domain-containing CAR T cells achieved better control of tumors over the rechallenge timepoints (**Figure 2b**). Collectively, our *in vitro* studies have identified three TAG72-CAR T cell lead candidates with potent but varying anti-tumor functional profiles, which we selected for further assessment of their *in vivo* preclinical therapeutic activity.

**Figure 2.**
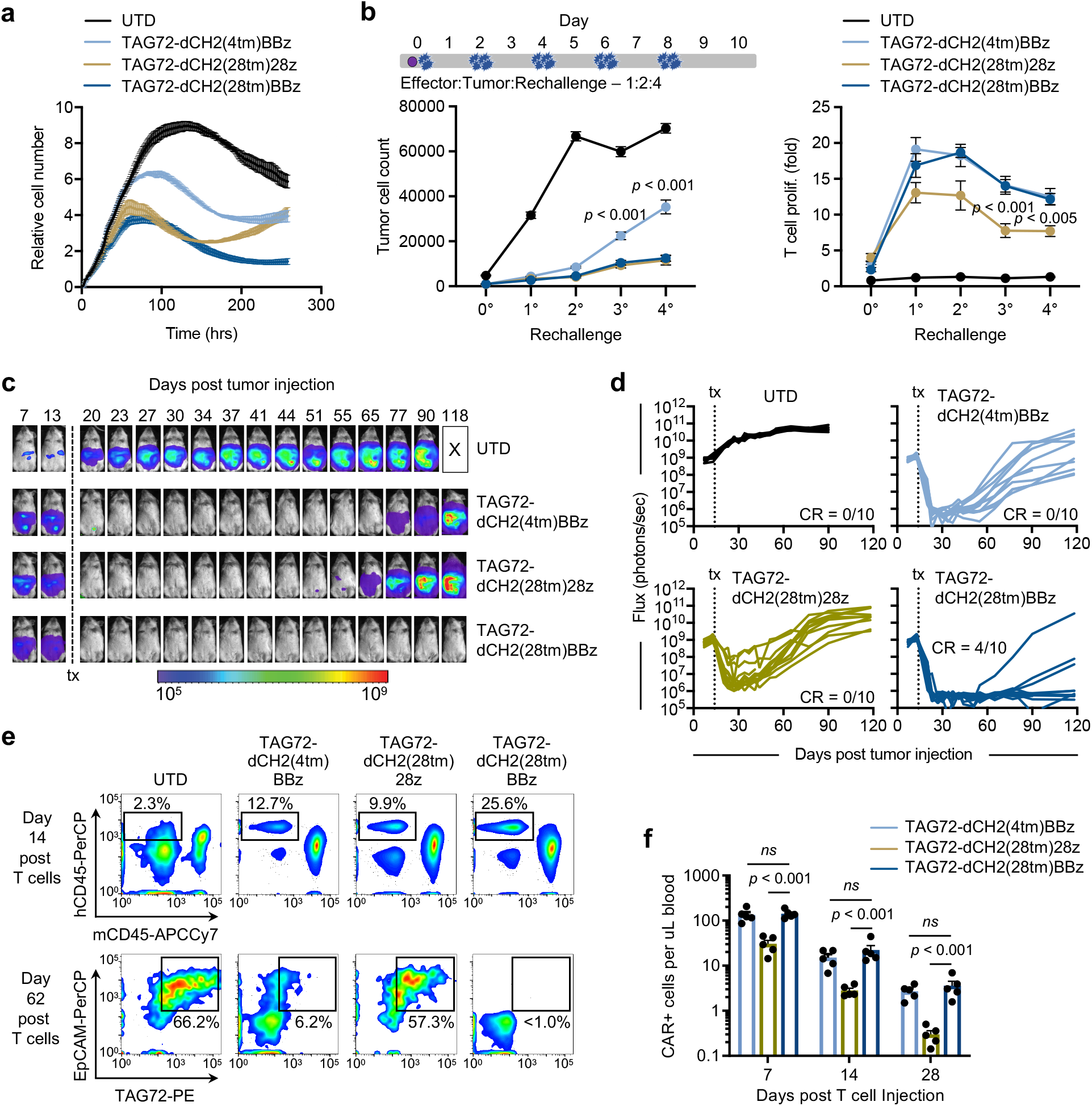
CD28 transmembrane domain in 4-1BB-based TAG72-CAR T cells induces durable therapeutic responses *in vitro* and *in vivo*. **(a)** TAG72-CAR T cell killing of OV90 cells measured by xCELLigence over 10 days (E:T = 1:20). **(b)** Schema of repetitive tumor cell challenge assay (top). TAG72-CAR T cells were co-cultured with OV90 cells (E:T = 1:2) and rechallenged with OV90 cells every two days. Remaining viable tumor cells and fold change in TAG72-CAR T cells were quantified as described in Materials and Methods prior to each tumor cell rechallenge. **(c)** Representative bioluminescent flux imaging of intraperitoneal (i.p.) OVCAR3(eGFP/ffluc) tumor-bearing female NSG mice treated i.p. with UTD or indicated TAG72-CAR T cells. **(d)** Quantification of flux (individual mice in each group) from treated OVCAR3 tumor-bearing mice. UTD (n = 8/group); TAG72-CAR T cells (n = 10/group). **(e)** Representative flow cytometric analysis of the frequency of human CD45+ (hCD45) and mouse CD45+ (mCD45) cells in the peritoneal cavity of tumor-bearing mice at day 14 post-treatment (top); human epithelial cell adhesion molecule+ (EpCAM) and tumor associated glycoprotein-72+ (TAG72) tumor cells in the i.p. cavity of tumor-bearing mice at day 62 post-treatment (bottom). **(f)** Quantification of TAG72-CAR T cells per uL of peripheral blood at 7, 14, and 28 days post-treatment. n = 5/group.

Anti-tumor efficacy of these three TAG72-CAR leads was evaluated using previously established human peritoneal ovarian tumor xenograft models^31^. At day 14 following intraperitoneal (i.p.) OVCAR3 tumor injection, mice were treated with untransduced (UTD), TAG72-dCH2(4tm)BBz, TAG72-dCH2(28tm)28z, or TAG72-dCH2(28tm)BBz CAR T cells (5.0 × 10^6^) by regional i.p. delivery. Dramatic anti-tumor responses were shown with all three CAR T cells, which was sustained for up to 6 weeks in treated mice (**Figure 2c**). However, we observed tumor recurrences after 6-8 weeks post-treatment in mice treated with TAG72-dCH2(4tm)BBz or TAG72-dCH2(28tm)28z CAR T cells, whereas TAG72-dCH2(28tm)BBz durably controlled tumors resulting in 4 out of 10 mice achieving complete therapeutic responses (**Figure 2d**).

To uncover the potential differences observed between the three TAG72-CAR T cell leads, we quantified CAR T cells in the peritoneal ascites and peripheral blood of mice following therapy. CAR T cells were seen in peritoneal ascites in treated mice, with TAG72-dCH2(28tm)BBz CAR T cells showing the greatest proportion of cells collected from the ascites 14 days after CAR T cell injection, compared to the other two CAR T cells (**Figure 2e top**). We additionally collected peritoneal ascites at 62 days post-treatment, at the time of tumor recurrences, and showed elimination of TAG72+ tumor cells in TAG72-dCH2(28tm)BBz CAR T cell-treated mice, while the presence of TAG72+ tumor cells remained with the other two CAR T cells (**Figure 2e bottom**). We observed similar numbers of 4-1BB costimulatory domain-containing CAR T cells in the blood of mice at three timepoints, higher than CD28 costimulatory domain-containing CAR T cells (**Figure 2f**). These data largely matched the T cell proliferation pattern in our *in vitro* functional assays. We further observed similar anti-tumor kinetics of the three CAR T cell lead candidates using a second, more aggressive, human OV90 peritoneal ovarian tumor xenograft model (**Supplemental Figure 4**).

To address potential safety concerns of this optimized TAG72-CAR containing a new anti-TAG72 scFv and CAR backbone, we evaluated the on- and off-target normal cell killing potential of TAG72-dCH2(28tm)BBz CAR T cells. Little to no TAG72 expression was observed across 10 primary human normal cells evaluated by flow cytometry, with minimal CD137+ T cell activation and cell killing by TAG72-CAR T cells, a finding that was consistent across five independent human CAR T cell donors (**Supplementary Figure 5**). In sum, the TAG72-dCH2(28tm)BBz CAR construct showed the most optimal anti-tumor functionality *in vitro* with safe, potent, and durable anti-tumor efficacy *in vivo*.

### IFN_γ_ signaling regulates the anti-tumor activity of CAR T cells

Accumulating evidence that IFNγ signaling impacts therapeutic activity of CAR T cells^39-41^, which has also been recently linked with CAR T cell killing capacity^42^. Along with our observation that TAG72-dCH2(28tm)BBz CAR T cells secreted the highest levels of IFNγ in our studies, we hypothesized that IFNγ signaling contributed to the superior anti-tumor activity of TAG72-dCH2(28tm)BBz CAR T cells. To test this hypothesis, we inhibited IFNγ signaling using an anti-IFNγR1 blocking antibody or enhanced IFNγ secretion with a human recombinant interleukin-12 (IL-12) in our extended *in vitro* co-culture tumor cell killing assay. Strikingly, we saw dose-dependent dampening of tumor cell killing with blockade of IFNγ signaling using OV90 tumors cells (**Figure 3a**) and OVCAR3 tumor cells (**Supplemental Figure 6**). We also observed dose-dependent enhancement of tumor cell killing with increasing concentrations of recombinant huIL-12, which resulted in the significant secretion of IFNγ by CAR T cells as determined by ELISA at the assay endpoint (**Figure 3b**). Interestingly, we observed only moderate impact on T cell proliferation following IFNγ blockade or addition of huIL-12 in this assay (**Figure 3c**). We further corroborated our findings using a recursive tumor cell killing assay, showing a requirement of IFNγ in promoting sustained anti-tumor activity *in vitro* (**Supplementary Figure 7**).

**Figure 3.**
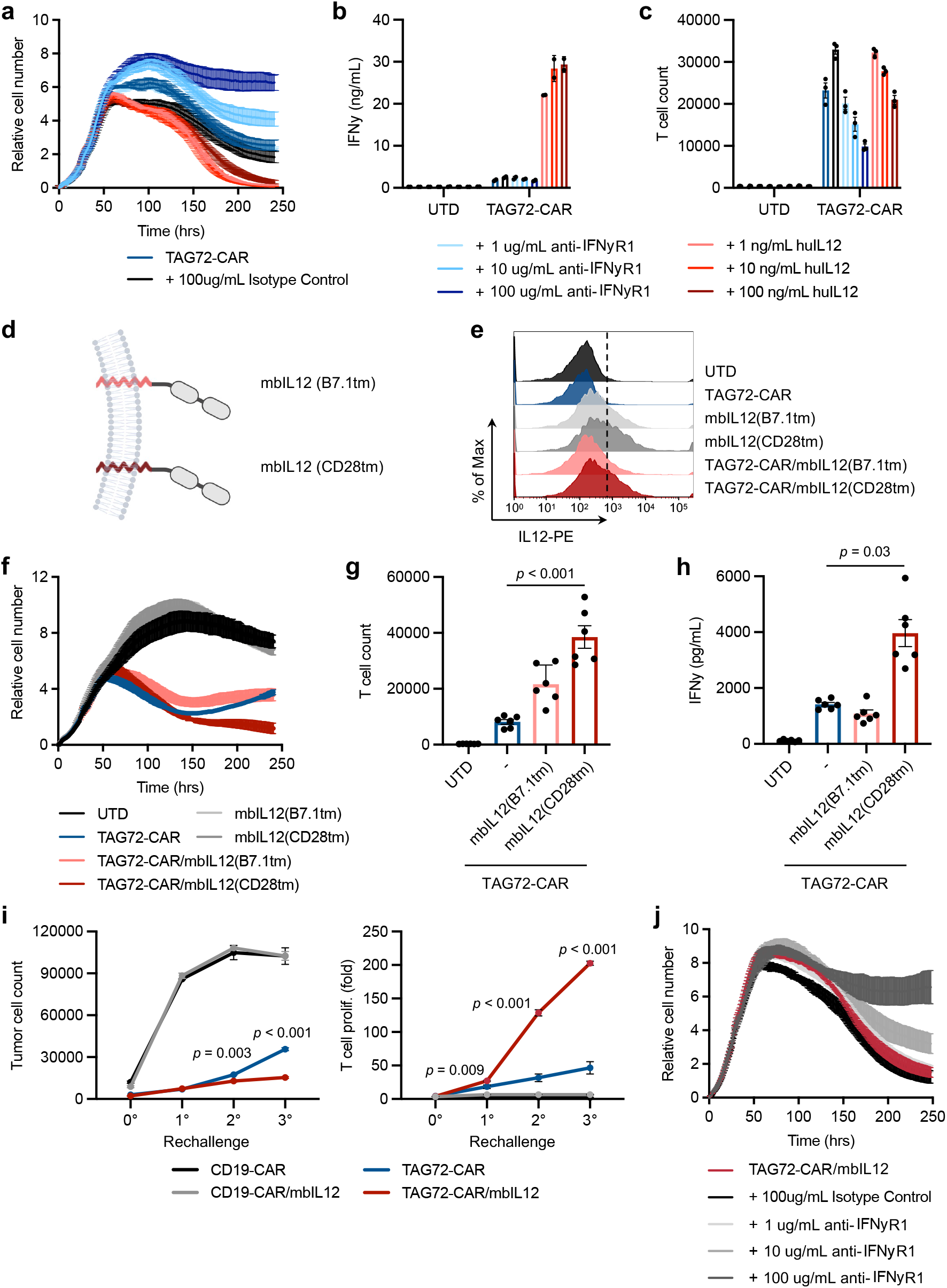
Membrane-bound IL-12 engineered TAG72-CAR T cells induce higher IFN_γ_,T cell expansion, and anti-tumor activity *in vitro*. **(a-c)** Tumor cell killing of OV90 cells by TAG72-CAR T cells (E:T = 1:20) with addition of varying concentrations of anti-IFNγR1 blocking antibody, isotype control, and recombinant human IL-12 cytokine measured by xCELLigence over 10 days (a). At day 10, IFNγ levels in supernatants were quantified by ELISA (b) and remaining T cell counts were analyzed by flow cytometry (c). **(d-h)** Diagram of the lentiviral construct with two versions of membrane-bound IL-12 (mbIL12) containing B7.1 or CD28 transmembrane domains (d). Flow cytometric analysis of IL-12 surface expression on indicated T cells (e). TAG72-CAR/mbIL12 T cell killing of OV90 cells (E:T = 1:20) measured by xCELLigence over 10 days (f). At day 10, remaining T cell counts were analyzed by flow cytometry (g) and IFNγ levels in supernatants were quantified by ELISA (h). **(i)** TAG72-CAR/mbIL12(CD28tm) T cells were co-cultured with OV90 cells (E:T = 1:3) and rechallenged with OV90 cells every two days. Remaining viable tumor cells and fold change in TAG72-CAR T cells were quantified as described in Materials and Methods prior to each tumor cell rechallenge.

### Engineered membrane-bound IL-12 signaling drives robust CAR T cell-mediated tumor cell killing and antigen-dependent T cell proliferation

Building on our *in vitro* findings that IL-12-induced IFNγ signaling is critical for CAR T cell anti-tumor activity, we engineered IL-12 into our TAG72-CAR T cells. Due to the potential off-target toxicity induced by secreted IL-12^43^, we aimed to spatially restrict IL-12’s effect by building membrane-bound IL-12 (mbIL12) constructs with varying transmembrane domains (**Figure 3d**). When we checked for their cell surface expression, we observed slightly enhanced expression of mbIL12(CD28tm) compared to mbIL12(B7.1tm), although both showed appreciable expression on TAG72-CAR T cells (**Figure 3e**). We then evaluated tumor cell killing activity of TAG72-CAR T cells expressing either of the two versions of mbIL12, and found that TAG72-CAR/mbIL12(CD28tm) T cells displayed the highest tumor cell killing activity, T cell expansion and IFNγ production (**Figure 3f-h**). Similar T cell functional benefits were observed using HER2-targeting and PSCA-targeting CAR T cells that were engineered with mbIL12(CD28tm) (**Supplemental Figure 8**). We further assessed activity of CAR T cells with mbIL12(CD28tm) in recursive tumor cell killing, in which we again observed enhanced tumor cell killing and CAR antigen-dependent T cell expansion over multiple rechallenge timepoints (**Figure 3i**). Importantly, no expansion or survival benefits were observed in the absence of CAR antigen stimulation (**Supplemental Figure 9**).

### Antigen-dependent IL-12 signaling in CAR T cells

We sought to better understand the signaling kinetics downstream of IL-12 and in the context of CAR T cell antigen stimulation. We first optimized an assay to confirm phosphorylated STAT4 (pSTAT4) downstream of recombinant human IL-12 (huIL12) in TAG72-CAR T cells. Interestingly, we observed that while pSTAT4 levels peaked at 1 hr and declined over the 24 hr timecourse with recombinant huIL12 alone, pSTAT4 was sustained over the 24 hr period in CAR T cells that were stimulated with plate-bound TAG72 antigen and huIL12 in combination **(Figure 4a-b)**. We also observed antigen-dependent induction of IL12RB2 expression, but not of constitutively-expressed IL12RB1, in CAR T cells (**Supplemental Figure 10**). We then compared the expression of cell surface mbIL12 on TAG72-CAR T cells that were rested in the absence of serum or exogenous cytokines prior to stimulation with plate-bound TAG72 antigen or control antigen. We observed that without stimulation, mbIL12 was detected at very low levels in engineered T cells. However, a significant and dose-dependent increase in cell surface expression of mbIL12 was seen in antigen-stimulated TAG72-CAR T cells, and in particular within CD137+ activated T cell subsets **(Figure 4c-e)**. Next, we interrogated pSTAT4 expression in TAG72-CAR/mbIL12 T cells in response to CAR stimulation. We observed the expected phosphorylation of STAT3 in response to TAG72 antigen in both TAG72-CAR and TAG72-CAR/mbIL12 T cells, which was only slightly activated by huIL12. However, CAR stimulation showed dose-dependent increases in pSTAT4 in TAG72-CAR/mbIL12 T cells compared to TAG72-CAR T cells alone **(Figure 4f)**. We further evaluated the potential for *trans* signaling in TAG72-CAR/mbIL12 T cells. We transduced HT1080 (TAG72-) cells with mbIL12 and co-cultured them with T cells in the presence of soluble CD3/CD28 stimulation. Increased pSTAT4 in T cells was observed when cultured with both HT1080-mbIL12 and soluble CD3/CD28 and not with HT1080-mbIL12 alone or HT1080-WT with CD3/CD28 **(Figure 4g)**. Collectively, these data suggest that mbIL12-engineered CAR T cells demonstrate improved *in vitro* anti-tumor activity and unexpectedly rely on CAR antigen stimulation, which we termed antigen-dependent IL-12 signaling.

**Figure 4.**
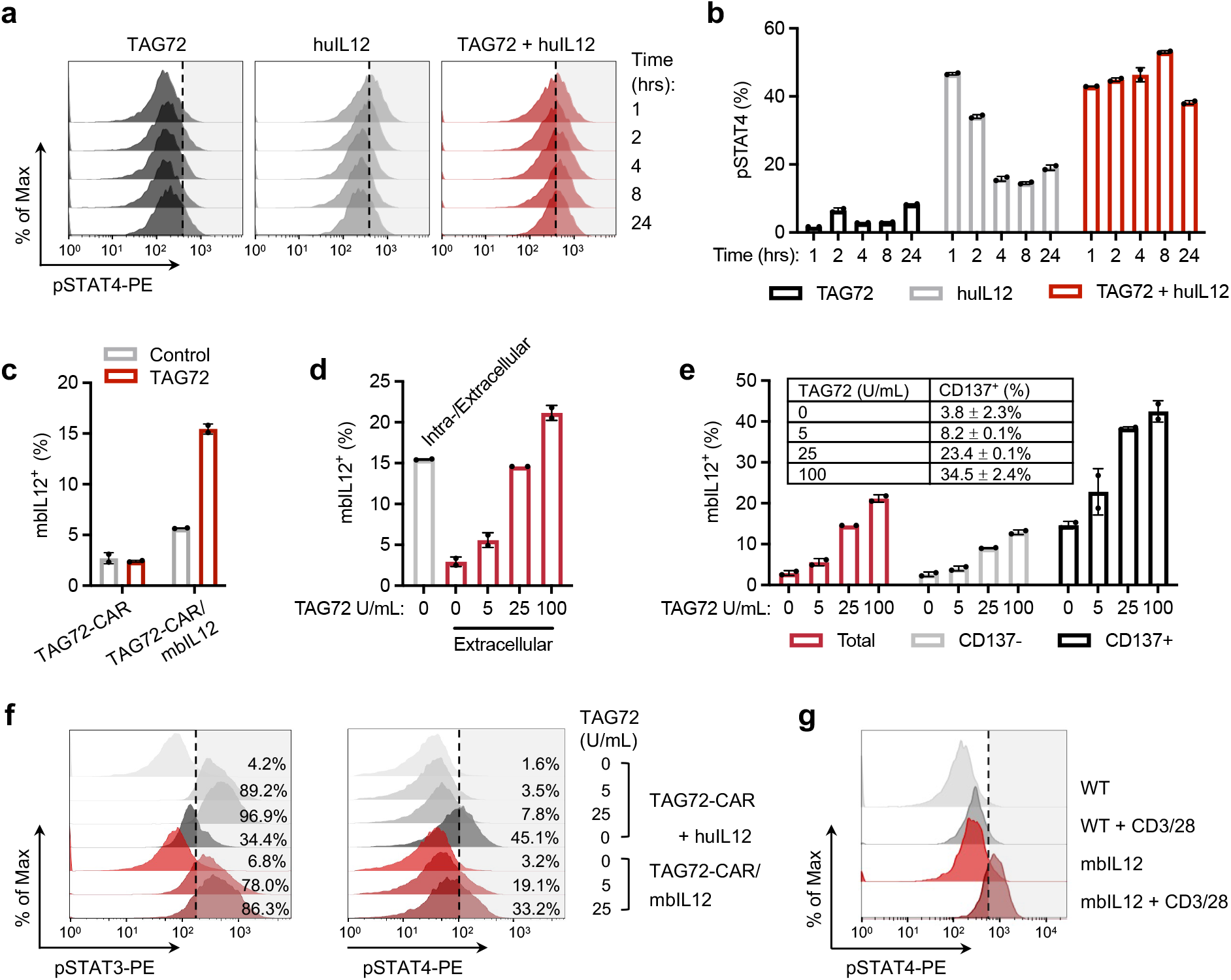
Antigen-dependent IL-12 signaling in TAG72-CAR T cells. **(a)** Representative intracellular flow cytometric analysis of phosphorylated STAT4 (pSTAT4, pY693) in response to TAG72 and/or recombinant huIL12 (10 ng/mL) at indicated timepoints. **(b)** Quantification of pSTAT4 in (a). **(c)** Flow cytometric analysis of surface expression of mbIL12 on TAG72-CAR T cells stimulated with plate-bound TAG72 (100 U/mL) or control antigen (PSCA, 2.5 ug/mL). **(d)** Flow cytometric analysis of surface or intracellular expression of mbIL12 in TAG72-CAR T cells stimulated with varying concentrations of plate-bound TAG72. **(e)** Flow cytometric analysis of surface expression of mbIL12 on total, or gated on CD137+ or CD137-populations, of TAG72-CAR T cells stimulated with plate-bound TAG72 (100 U/mL). Table inset: CD137+ expression on TAG72-CAR T cells following stimulation with varying concentrations of plate-bound TAG72. **(f)** Intracellular flow cytometric analysis of phosphorylated STAT3 (pSTAT3, pY705) (left) and pSTAT4 (right) in TAG72-CAR and TAG72-CAR/mbIL12 T cells stimulated with varying concentrations of plate-bound TAG72 or recombinant huIL12 (10 ng/mL). **(g)** Intracellular flow cytometric analysis of pSTAT4 in TAG72-CAR T cells co-cultured with HT1080 (TAG72-) cells transduced with mbIL12. Cells were stimulated with Immunocult CD3/CD28 per manufacturer recommendation. Cells were gated on CAR T cells and evaluated for pSTAT4.

### Superior anti-tumor activity by antigen-dependent IL-12 signaling in CAR T cells

We next evaluated the therapeutic potential of TAG72-CAR T cells with antigen-dependent IL-12 signaling. Using the i.p. OVCAR3 tumor xenograft model, mice treated with TAG72-CAR/mbIL12 T cells sustained more durable anti-tumor responses as compared to TAG72-CAR T cells, achieving a greater incidence of durable complete responses (**Figure 5a-b**). Importantly, tumor-bearing mice treated with CD19-CAR T cells and CD19-CAR/mbIL12 T cells showed little differences in therapy, supporting the antigen-dependent nature of mbIL12. We observed a higher frequency of hCD45+ cells in the peritoneal ascites of mice treated with TAG72-CAR/mbIL12 T cells as compared to mice treated with TAG72-CAR T cells alone at 2 and 4 weeks post-treatment (**Figure 5c-d**). We also observed significantly higher and sustained levels of TAG72-CAR/mbIL12 T cells in peripheral blood as compared with TAG72-CAR T cells alone (**Figure 5d**). We replicated these findings using the more challenging i.p. OV90 tumor xenograft model, in which the differences in therapy and T cell persistence between TAG72-CAR and TAG72-CAR/mbIL12 T cells were even more pronounced (**Figure 5e-f**).

**Figure 5.**
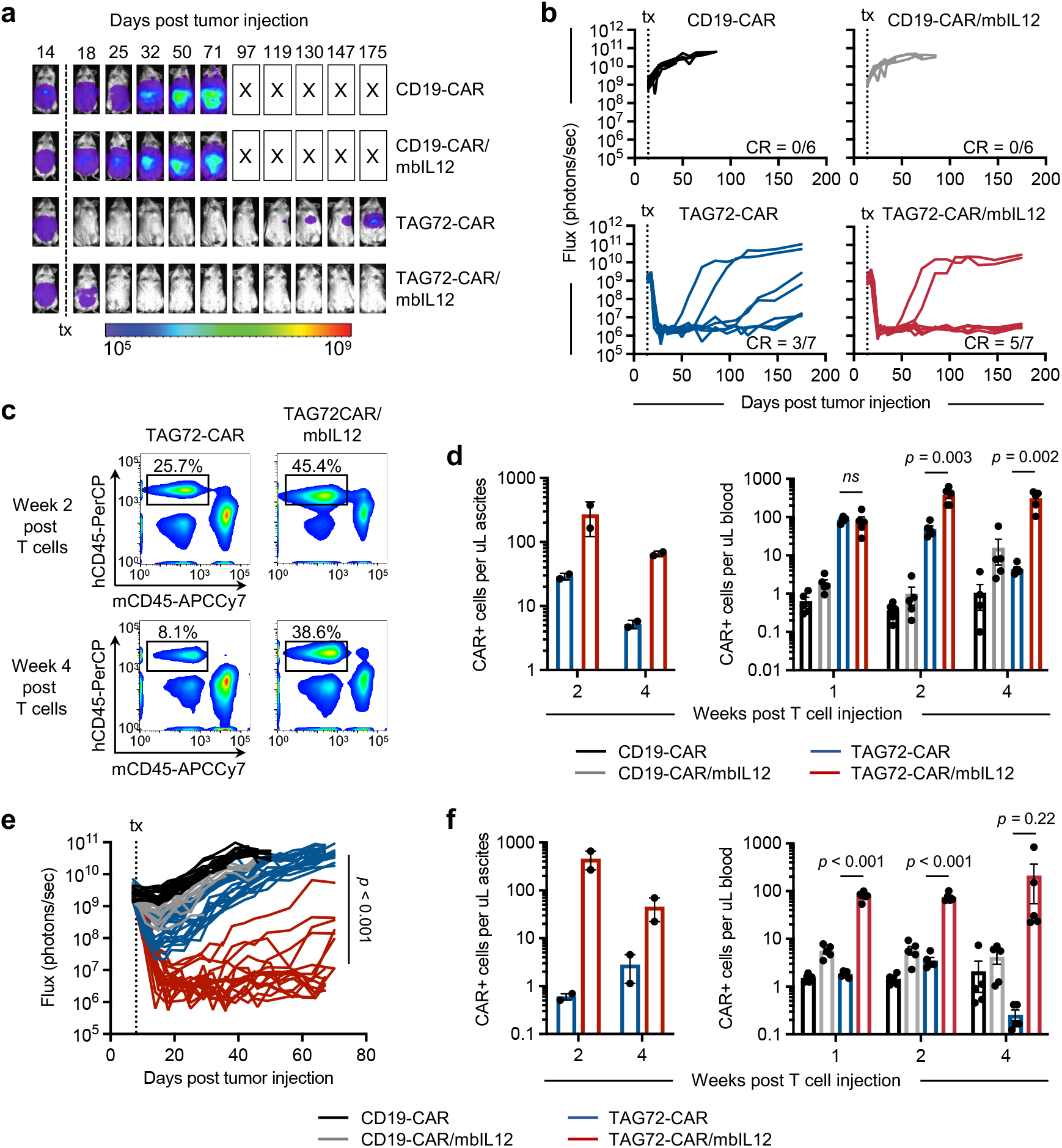
Locoregional intraperitoneal delivery of TAG72-CAR/mbIL12 T cells reduces tumor burden and increases regional and systemic CAR T cell persistence *in vivo*. **(a)** Representative bioluminescent flux imaging of i.p. OVCAR3(eGFP/ffluc) tumor-bearing mice treated i.p. with CD19-CAR, CD19-CAR/mbIL12, TAG72-CAR or TAG72-CAR/mbIL12 T cells. **(b)** Quantification of flux (individual mice per group) from mice treated i.p. with CD19-CAR T cells (n = 6/group), CD19-CAR/mbIL12 T cells (n = 6/group), TAG72-CAR T cells (n = 7/group) and TAG72-CAR/mbIL12 T cells (n = 7/group). **(c)** Representative flow cytometric analysis of human CD45+ (hCD45) and mouse CD45+ (mCD45) cells in the peritoneal cavity of tumor-bearing mice at week 2 (top) or week 4 (bottom) post-treatment. **(d)** Quantification of TAG72-CAR T cells per uL of peritoneal ascites (left) at weeks 2 and 4 post-treatment. n = 2 per group. Quantification of TAG72-CAR T cells per uL of peripheral blood (right) at weeks 1, 2, and 4 post-treatment. n = 5/group. **(e)** Quantification of flux (individual mice per group) from OV90(eGFP/ffluc) tumor-bearing mice treated i.p. with CD19-CAR T cells (n = 5/group), CD19-CAR/mbIL12 T cells (n = 5/group), TAG72-CAR T cells (n = 9-10/group) and TAG72-CAR/mbIL12 T cells (n = 9 - 10/group). Combined data are from two independent studies. **(f)** Quantification of TAG72-CAR T cells per uL of peritoneal ascites (left) at weeks 2 and 4 post-treatment. n = 2 per group. Quantification of TAG72-CAR T cells per uL of peripheral blood (right) at weeks 1, 2, and 4 post-treatment. n = 5/group.

### Improved systemic disease targeting by CAR T cells with antigen-dependent IL-12 signaling

One prevailing argument against the locoregional administration of CAR T cells is their potential spatial confinement, thereby preventing systemic therapy in patients with widespread metastatic disease^44^. However, our data suggest that regional intraperitoneally-administered mbIL12-engineered CAR T cells may have a greater capacity to target disease outside of the peritoneum. To test this, we established a xenograft OV90 tumor model with both a regional i.p. and systemic s.c. tumor in the same mouse. OV90 (ffluc-negative) tumor cells were injected subcutaneously (s.c.) to track with calipers measurement and OV90 (eGFP/ffluc-expressing) tumor cells were i.p. injected and tracked with bioluminescent flux imaging (**Figure 6a**). As we observed in previous experiments, i.p. anti-tumor responses were greater in mice regionally treated with TAG72-CAR/mbIL12 T cells as compared to TAG72-CAR T cells alone (**Figure 6b-c**). While s.c. tumors initially regressed similarly in both treatment groups, all tumors recurred following TAG72-CAR T cell treatment alone, whereas s.c. tumors were completely eradicated in all mice followingTAG72-CAR/mbIL12 T cell treatment (**Figure 6d**). We corroborated this phenomenon by again observing higher levels of CAR T cells in the peripheral blood and peritoneal ascites of mice treated with TAG72-CAR/mbIL12 T cells (**Figure 6e-f**). Furthermore, immunohistochemistry (IHC) analysis of s.c. tumors at day 12 post treatment demonstrated significantly greater infiltration of TAG72-CAR/mbIL12 T cells as compared with TAG72-CAR T cells alone (**Figure 6g**). Overall, these data support locoregional delivery of CAR T cells with engineered antigen-dependent IL-12 signaling in durable targeting of both regional and systemic disease.

**Figure 6.**
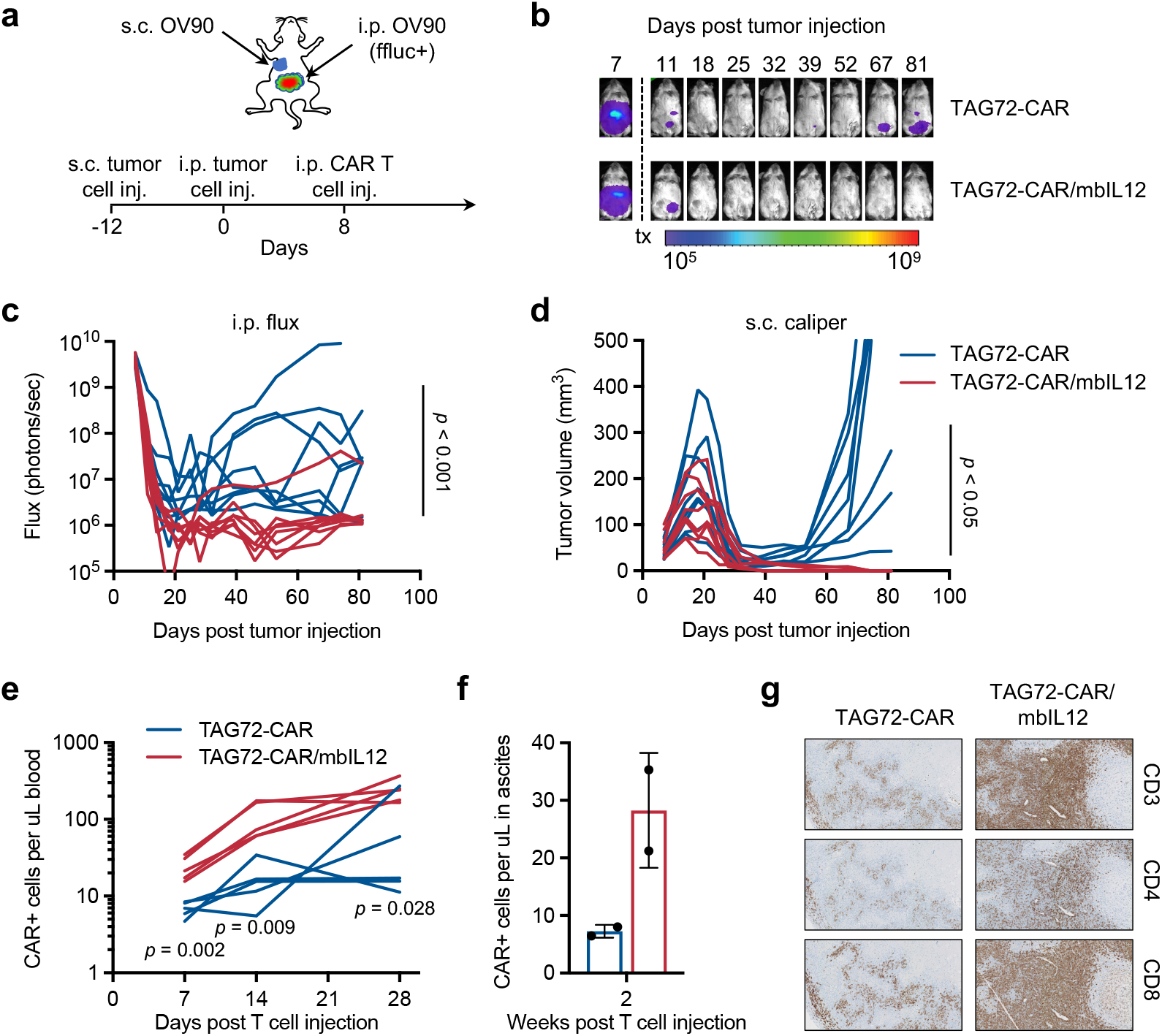
Locoregional intraperitoneal delivery of TAG72-CAR/mbIL12 T cells eradicate subcutaneous tumors in dual tumor-bearing mice. **(a)** Schematic for subcutaneous (s.c.) and i.p. OV90 dual tumor model and treatment. **(b)** Representative bioluminescent flux imaging of dual tumor-bearing mice treated i.p. with TAG72-CAR or TAG72-CAR/mbIL12 T cells. **(c)** Quantification of flux (individual mice per group) from mice treated i.p. with TAG72-CAR T cells (n = 8/group) and TAG72-CAR/mbIL12 T cells (n = 8/group). **(d)** Quantification of subcutaneous tumor volume (individual mice per group) from mice treated i.p. with TAG72-CAR T cells. **(e)** Quantification of TAG72-CAR T cells per uL of peripheral blood at days 7, 14, 21, and 28 post-treatment. n = 5/group. **(f)** Quantification of TAG72-CAR T cells per uL of peritoneal ascites at week 2 post-treatment. n = 2/group. **(g)** Representative immunohistochemistry analysis of CD3+ T cells in s.c. tumors at day 12 post-treatment.

## Discussion

Our findings address major challenges associated with engineering safe and effective CAR T cell strategies for solid tumors. Here, we fully optimized TAG72-specific CARs by varying the antigen-binding scFv, extracellular spacer, transmembrane, and intracellular costimulatory domains, and also validated the lead candidate using increasingly challenging *in vitro* studies and *in vivo* xenograft models of ovarian cancer. We are now underway in evaluating safety, feasibility, and bioactivity of regionally-delivered TAG72-CAR T cells in patients with advanced ovarian cancer in an open phase 1 trial (NCT05225363). Perhaps the most critical modification to the CAR was incorporation of the CD28 transmembrane domain to the 4-1BB costimulatory domain, which allowed for durable anti-tumor activity and improved T cell expansion in recursive tumor cell killing assays. Further, we identified IL-12-regulated IFNγ signaling in driving cytolytic activity and expansion/persistence of CAR T cells *in vitro* and *in vivo*. Unexpectedly and perhaps most desirable, we observed that cell surface expression and signaling of our membrane-bound IL-12 molecule was tightly controlled by antigen-dependent CAR activation.

Based on our own findings as well as emerging data in the CAR T cell field, we reasoned that our clinical lead TAG72-CAR T cell’s superior performance against solid tumors is due to its ability to secrete high levels of IFNγ and enhance tumor cell killing. This notion was supported by demonstrating dampened TAG72-CAR T cell activity with blockade of IFNγR1 signaling, which is supported by a recent study by Larson *et al*^42^. consequently, we now show enhanced CAR T cell cytotoxicity with exogenous IL-12, a cytokine that canonically induces IFNγ secretion in T cells^30,45,46^. We further engineered CAR T cells targeting TAG72, HER2, or PSCA, with a membrane-bound IL-12 molecule and showed similar improvements in anti-tumor activity. While we did not perform studies in syngeneic models, research by other groups have demonstrated impressive impact of IL-12 signaling in the immunosuppressive tumor microenvironment, with effects that also likely influence tumor vasculature through IFNγ signaling to promote anti-tumor responses^30,41^. In support of these dynamic IL-12-driven effects, which warrants further investigation, we showed that mbIL12 signaling in T cells can also signal in *trans* with the capacity to impact other cell types within the solid tumor microenvironment. We believe these data have broad applicability to other CAR T cell strategies and disease settings.

Important to the CAR T cell regional administration approach we used in this study and in the clinical design of our upcoming phase 1 trial, we observed greater CAR T cell persistence in the peritoneal ascites as well as in peripheral blood of treated mice with CAR T cells further engineered with mbIL12. The durable persistence of TAG72-CAR T cells in the periphery led us to hypothesize that mbIL12 signaling in CAR T cells could solve one of the major hurdles facing local or regional delivery approaches, being the limited biodistribution and targeting of widespread metastatic disease. Indeed, our finding that regionally-delivered mbIL12-endowed CAR T cells better controlled regional as well as systemic disease compared to CAR T cells alone is a strong contender to resolving this issue. One major shortcoming in our study is that we did not evaluate other models of regional to systemic distribution of CAR T cells, such as the intraventricular delivery of CAR T cells for the treatment of either primary brain tumors or multifocal brain metastases^23^. However, we reason that this phenomenon is likely applicable beyond regional intraperitoneal delivery.

Conditional or inducible gene engineering in CAR T cells is a major advancement to the field, with the potential to greatly improve safety and overall efficacy of solid tumor-directed CAR T cells^3,47^. Various spatially and temporally activated gene promoters which drive expression of CAR or other genes have been widely explored, ranging from heat-, light-, and hypoxia-inducible promoters, as well as synthetic Notch and drug-induced circuits^48-53^. While many of these approaches have not yet been investigated clinically, previous attempts to regulate gene expression by linking to conditionally-activated promoters, including the NFAT promoter^54^, have been clinically tested. Unfortunately, recent attempts at regulating expression of a secretable IL-12 under an inducible NFAT promoter in an adoptively transferred T cell therapy still resulted in unwanted toxicities due to systemic distribution of the cytokine^43^. To solve this issue, many groups have evaluated membrane-bound or -tethered approaches to limit distribution of immune-modulating cytokines, including IL-12 and IL-15^55-58^. Here, we tethered the immune-modulating IL-12 to the surface of CAR T cells using an optimized CD28 transmembrane domain and discovered that its sustained cell surface expression and downstream signaling effects on CAR T cells were antigen-dependent. The precise mechanisms underlying this conditional mbIL12 signaling is under investigation by our group, and may be a viable approach to broadly regulate T cell cytokine signaling including IL-2, IL-7, IL-15, and others.

In summary, this work highlights advantages of comprehensively optimizing CAR functionality by varying different regions on the CAR molecule combined with systematically testing functional differences through challenging *in vitro* and *in vivo* preclinical modeling. We linked the enhancement of solid tumor killing by CAR T cells with the increased presence of IFNγ signaling, which we achieved by engineering antigen-dependent mbIL12 on CAR T cells. This addition not only improved regional tumor control but also enhanced systemic expansion and persistence of TAG72-CAR T cells to promote eradication of systemic disease in our model system. Our findings have the potential to broadly improve CAR T cell therapeutic responses against multi-metastatic diseases.

## Materials and Methods

### Cell lines

The epithelial ovarian cancer line OVCAR3 (ATCC HTB-161) was cultured in RPMI-1640 (Lonza) containing 20% fetal bovine serum (FBS, Hyclone) and 1X antibiotic-antimycotic (1X AA, Gibco) (complete RPMI). The epithelial ovarian cancer line derived from metastatic ascites OV90 (CRL-11732) was cultured in a 1:1 mixture of MCDB 105 medium (Sigma) and Medium 199 (Thermo) adjusted to pH of 7.0 with sodium hydroxide (Sigma) and final 20% FBS and 1X AA. The epithelial ovarian cancer line OVCAR8 was a generous gift from Dr. Carlotta Glackin at City of Hope and was cultured in complete RPMI-1640. All cells were cultured at 37°C with 5% CO_2_. Human primary cell lines were obtained from Cell Biologics (Human Primary Colonic Epithelial Cells H-6047, Human Primary Esophageal Epithelial Cells H-6046, Human Primary Kidney Epithelial Cells H-6034, Human Primary Ovarian Epithelial Cells H-6036, Human Primary Pancreatic Epithelial Cells H-6037, Human Primary Proximal Tubular Epithelial Cells H-6015, Human Primary Small Intestine Epithelial Cells H-6051, Human Primary Stomach Epithelial Cells H-6039), Promocell (Human Cardiac Myocytes C-12810), and Lonza (Human Bronchial Epithelial Cells CC-2541) and cultured according to vendor’s specifications.

### DNA constructs, tumor lentiviral transduction, and retrovirus production

Tumor cells were engineered to express enhanced green fluorescent protein and firefly luciferase (eGFP/*ffluc*) by transduction with epHIV7 lentivirus carrying the eGFP/*ffluc* fusion under the control of the EF1α promoter as described previously^23^. The humanized scFv sequence used in the CAR construct was obtained from a monoclonal antibody clone huCC49 that targets TAG72^31^. The extracellular spacer domain included the 129-amino acid middle-length CH2-deleted version (ΔCH2) of the IgG4 Fc spacer^31^. The intracellular co-stimulatory signaling domain contained was a 4-1BB with a CD4 transmembrane domain. The CD3ζ cytolytic domain was previously described^31^. Variations in extracellular spacer domains, transmembrane domains, and intracellular co-stimulatory signaling domains were described previously^11,23,38^. The CAR sequence was separated from a truncated CD19 gene (CD19t) by a T2A ribosomal skip sequence, and cloned in an epHIV7 lentiviral backbone under the control of the EF1α promoter. The PSCA-BBζ CAR construct was described previously^38^. The membrane-bound IL-12 (mbIL12) construct was generated using the p35 and p40 genes (p35, NC_000003.12; p40, NC_000005.10) separated by a G4S spacer, and linked to either the B7.1 or CD28 transmembrane domain. Lentivirus was generated as previously described^38^. Lentiviral titers were quantified using HT1080 cells based on CD19t or IL-12 cell surface expression using flow cytometry.

### T cell isolation, lentiviral transduction, and *ex vivo* expansion

Leukapheresis products were obtained from consented research participants (healthy donors) under protocols approved by the City of Hope Internal Review Board (IRB), and enriched for T cells as previously described^38,59^. T cell activation and transduction was performed as described previously^38^. Where indicated, we performed a second lentiviral transduction followed 24 hr after the first transduction. Cells were then *ex vivo* manufactured, enriched for CAR, and frozen as described previously^38^. Purity and cell surface phenotype of CAR T cells were analyzed by flow cytometry using antibodies and methods as described below.

### Flow cytometry

For flow cytometric analysis, cells were resuspended in FACS buffer (Hank’s balanced salt solution without Ca^2+^, Mg^2+^, or phenol red (HBSS^−/−^, Life Technologies) containing 2% FBS and 1 × AA). Cells were incubated with primary antibodies for 30 min at 4°C in the dark. For secondary staining, cells were washed twice prior to 30 min incubation at 4°C in the dark with either Brilliant Violet 510 (BV510), fluorescein isothiocyanate (FITC), phycoerythrin (PE), peridinin chlorophyll protein complex (PerCP), PerCP-Cy5.5, PE-Cy7, allophycocyanin (APC), or APC-Cy7 (or APC-eFluor780)-conjugated antibodies. Antibodies against CD3 (BD Biosciences, Clone: SK7), CD4 (BD Biosciences, Clone: SK3), CD8 (BD Bosciences, Clone: SK1), CD19 (BD Biosciences, Clone: SJ25C1), mouse CD45 (BioLegend, Clone: 30-F11), CD45 (BD Biosciences, Clone: 2D1), CD69 (BD Biosciences, Clone: L78), CD137 (BD Biosciences, Clone: 4B4-1), Ep-CAM/CD326 (BioLegend, Clone: 9C4), biotinylated Protein L (GenScript USA) (25), TAG72 (Clone, muCC49), Donkey Anti-Rabbit Ig (Invitrogen), Goat Anti-Mouse Ig (BD Biosciences), and streptavidin (BD Biosciences) were used. Cell viability was determined using 4′, 6-diamidino-2-phenylindole (DAPI, Sigma). Flow cytometry was performed on a MACSQuant Analyzer 10 (Miltenyi Biotec), and the data was analyzed with FlowJo software (v10, TreeStar).

For intracellular flow cytometry, CAR T cells were thawed and rested in IL-2 (50 U/mL) & IL-15 (0.5 ng/mL) overnight at 1 × 10^6^ cells/mL. On the following day, CAR T cells were washed twice in 1x PBS and suspended at 1 × 10^6^ cells/mL in media without serum or cytokines. 1 × 10^5^ cells were plated per well in a 96-well plate to rest overnight. The next day, cells were stimulated with either soluble cytokine [IL-2 (50 U/mL), IL-15 (0.5 ng/mL), IL-12 (10 ng/mL)] or transferred to a high-binding 96-well plate pre-coated with indicated amounts of control or TAG72 antigen (BioRad). Reagents and buffers for flow cytometry processing were pre-chilled on ice unless otherwise stated. Following antigen stimulation, cells were washed with FACS buffer (supplemented with 0.1% sodium azide) and then fixed in pre-warmed 1x BD Phosflow Lyse/Fix buffer (558049) at 37°C for 10 minutes. Cells were then washed with FACS buffer and if required, stained with the extracellular antibodies on ice for 30 minutes in the dark. Stained cells were washed and suspended in pre-chilled (−20°C) BD Perm Buffer III (558050) and kept on ice for 30 minutes. Following a wash, cells were suspended in human FC block (Miltenyi Biotec Inc., FLP3330) and kept on ice for 30 minutes, washed and stained with intracellular antibodies: PE-pSTAT3, PE-pSTAT4, PE-pSTAT5 (Biolegend). Data was acquired on a MACSQuant Analyzer 16 cytometer (Miltenyi) and analyzed with FlowJo v10.8.

### *In vitro* tumor killing and T cell functional assays

For tumor cell killing assays, CAR T cells and tumor targets were co-cultured at indicated effector:tumor (E:T) ratios in complete X-VIVO without cytokines in 96-well plates for the indicated time points and analyzed by flow cytometry as described above. Tumor cells were plated overnight prior to addition of T cells. Tumor cell killing by CAR T cells was calculated by comparing CD45-negative DAPI-negative (viable) cell counts relative targets co-cultured with untransduced (UTD) T cells. For xCELLigence tumor cell killing assays, CAR T cells and tumor targets were co-cultured at indicated effector:tumor (E:T) ratios in complete X-VIVO without cytokines in 96-well plates for up to 10 days and analyzed by flow cytometry as described above.

For T cell activation assays, CAR T cells and tumor targets were co-cultured at the indicated E:T ratios in complete X-VIVO without cytokines in 96-well plates for the indicated time points and analyzed by flow cytometry for indicated markers of T cell activation. For T cell activation assays on plate-bound antigen, purified soluble TAG72 antigen (BioRad) was plated in duplicate at indicated TAG72 units overnight at 4°C in 1X PBS in 96-well flat bottom high-affinity plates (Corning). Using a Bradford protein assay, the 20,000 units/mL stock solution of soluble TAG72 antigen was determined to be approximately 1.234 mg/mL of total protein. A designated number of TAG72-CAR T cells were then added in a fixed volume of 100 uL to each well and incubated for indicated times prior to collection of cells for analysis of activation markers (CD69, CD137) by flow cytometry. Supernatants were also collected for analysis of cytokine production. For T cell survival assays, T cells were plated at 1 × 10^6^ cells/mL in X-VIVO 10% FBS with or without cytokines and counted every two days. Cell concentration was adjusted to 1 × 10^6^ cells/mL with fresh media following each count day.

### ELISA cytokine assays

Supernatants from tumor cell killing assays or CAR T cell activation assays on plate-bound TAG72 antigen were collected at indicated times and frozen at -20°C for further use.

Supernatants were then analyzed for secreted human IFNγ and IL-2 according to the Human IFNγ and IL-2 ELISA Ready-SET-GO!^®^; ELISA kit manufacturer’s protocol, respectively. Plates were read at 450 nm using a Wallac Victor3 1420 Counter (Perkin-Elmer) and the Wallac 1420 Workstation software.

### Western blotting analysis

Cell pellets were thawed on ice. After thaw, cell pellets were resuspended in RIPA buffer consisting of 25mM Tris-HCl (pH 8.5), 150mM NaCl, 1mM EDTA (pH 8.0), 1%(v/v) NP-40 substitute, 0.5%(w/v) Sodium Deoxycholate, 0.1%(w/v) SDS, 10mM NaF, 1mM NaOV, 10mM β-glycerophosphate, and 1x of Halt Protease and Phosphotase Inhibitor Cocktail (Thermo Scientific). Lysates were incubated on ice for 30 min then centrifuged at 17,200*g* for 20 minutes at 4°C. Lysate supernatant was transferred to a new tube and anylized for total protein concentration by Braford protein assay. Laemmli sample buffer (BioRad) containing DTT (Sigma Aldrich) was added to propotional quantities of total protein and samples were poiled at 95°C for 5 minutes. Protein was sepearted on a 7.5% Criterion TGX Precast Midi Protein Gel (BioRad) using the Criterion Cell (BioRad) and transferred to 0.2um nitrocellulose blotting membrane (Genesee) in Tris-Glycine Transfer Buffer (Thermo Scientific) using the Trans-Blot Turbo Electrophoretic Transfer Cell (BioRad). Membranes were washed in deionized water, incubated in Ponceau S solution (Sigma Aldrich) to confirm protein transfer, and then washed in Tris-buffered saline containing 0.05% Tween20 (Sigma Aldrich) (TBST) for 1 minute. Membranes were then blocked for 1 hour at room temperature in blocking buffer containing 5% PhosphoBLOCKER blocking reagent (Cell Biolabs) in TBST. After blocking, membranes were transferred to blocking buffer containing primary antibodies and incubated overnight at 4°C. All primary antibodies were sourced from Cell Signaling Technology and included actin (1:2000), p44/42 MAPK (ERK1/2) (1:1000), pp44/42 MAPK (pERK1/2) (1:1000), SLP76 (1:1000), pSLP76 (1:1000), PLCγ1 (1:1000), and pPLCγ1 (1:1000). Membranes were washed in TBST and then incubated for 45 min at room temperature in blocking buffer containing either anti-rabbit or anti-mouse HRP-linked secondary antibody (Cell Signaling Technology). Membranes were washed in TBST and imaged on the ChemiDoc Imaging System using SuperSignal chemiluminescent substrate (Thermo Scientific).

### *In vivo* studies

All animal experiments were performed under protocols approved by the City of Hope Institutional Animal Care and Use Committee. For *in vivo* intraperitoneal (i.p.) tumor studies, OVCAR3 and OV90 cells (5.0 × 10^6^) were prepared in a final volume of 500 uL HBSS^−/−^ and engrafted in >6 weeks old female NSG mice by i.p. injection. For subcutaneous (s.c.) tumor studies, OV90 cells (5 × 10^6^) were prepared in a final volume of 100 uL HBSS^−/−^ and injected under the skin of the abdomen of 6–8 weeks old female NSG mice. Tumor growth was monitored at least once a week via non-invasive bioluminescence imaging (Xenogen, LagoX) and flux signals were analyzed with Living Image software (Xenogen). For imaging, mice were i.p. injected with 150 uL D-luciferin potassium salt (Perkin Elmer) suspended in PBS at 4.29 mg/mouse. At day 8 for OV90 and day 14 for OVCAR3, mice were i.p. treated with indicated T cells (5 × 10^6^) in 500 uL final volume. Humane endpoints were used in determining survival. Mice were euthanized upon signs of distress such as a distended belly due to peritoneal ascites, labored or difficulty breathing, apparent weight loss, impaired mobility, or evidence of being moribund. At pre-determined time points or at moribund status, mice were euthanized and tissues and/or peritoneal ascites were harvested and processed for flow cytometry and immunohistochemistry as described above.

Peripheral blood was collected from isoflurane-anesthetized mice by retro-orbital (RO) bleed through heparinized capillary tubes (Chase Scientific) into polystyrene tubes containing a heparin/PBS solution (1000 units/mL, Sagent Pharmaceuticals). Total volume of each RO blood draw (approximately 120 uL/mouse) was recorded. Red blood cells (RBCs) were lysed with 1X Red Cell Lysis Buffer (Sigma) according to manufacturer’s protocol and then washed, stained, and analyzed by flow cytometry as described above. Cells from peritoneal ascites were collected from euthanized mice by injecting 5 mL cold 1X PBS into the i.p. cavity, which was drawn up via syringe and stored on ice until further processing. RBC-depleted ascites cells were washed, stained, and analyzed by flow cytometry using antibodies and methods as described above.

### Immunohistochemistry

Tumor tissue was fixed for up to 3 days in 4% paraformaldehyde (4% PFA, Boston BioProducts) and stored in 70% ethanol until further processing. Immunohistochemistry was performed by the Pathology Core at City of Hope. Briefly, paraffin-embedded sections (10 um) were stained with hematoxylin & eosin (H&E, Sigma-Aldrich), mouse anti-human CD3 (DAKO), mouse anti-human CD4 (DAKO), mouse anti-human CD8 (DAKO), and mouse anti-human TAG72 (AB16838, Abcam). Images were obtained using the Nanozoomer 2.0HT digital slide scanner and the associated NDP.view2 software (Hamamatzu).

## Statistical anslysis

Data are presented as means ± standard error mean (SEM), unless otherwise stated. Statistical comparisons between groups were performed using the unpaired two-tailed Student’s *t* test to calculate *p* value, unless otherwise stated.

## Supplemental Information

Supplemental figures and legends are available and included as separate document.

## Acknowledgments

We thank the staff members of the following cores at the Beckman Research Institute at City of Hope Comprehensive Cancer Center: Animal Facility, Pathology, and Small Animal Imaging for their excellent technical assistance.

## Author contributions

SJP, along with EHL and JPM provided conception and construction of the study. SJP, LDW, SJF, LRR, MC, YW, JPM, CC, and EHL provided design of experimental procedures, data analysis, and/or interpretation. EHL, CC, JPM, DG, AKP, JY, LS, LNA, GD, BG, and WCC performed experiments. SJP, EHL, and CC wrote the manuscript. CM assisted in writing/editing the manuscript. SJP supervised the study. All authors reviewed the manuscript.

