## Supplemental Figures for "Antigen-dependent IL-12 signaling in CAR T cells promotes regional to systemic disease targeting"

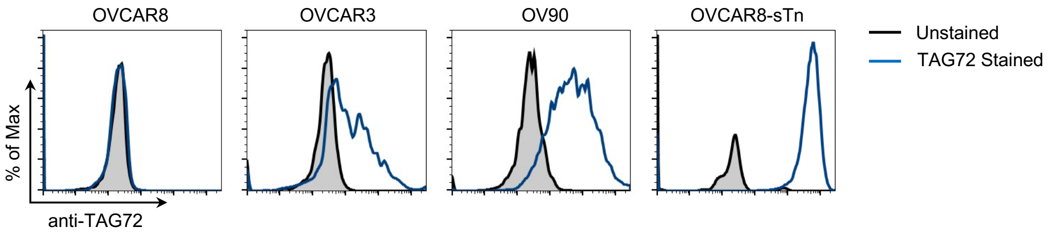


**Supplemental Figure 1. TAG72 expression on ovarian tumor cell lines.** Flow cytometric analysis of TAG72 expression on human OVCAR8 (TAG72-negative), human OVCAR3 (TAG72-positive), human OV90 (TAG72-positive), human OVCAR8-sTn (TAG72-positive) tumor cell lines.

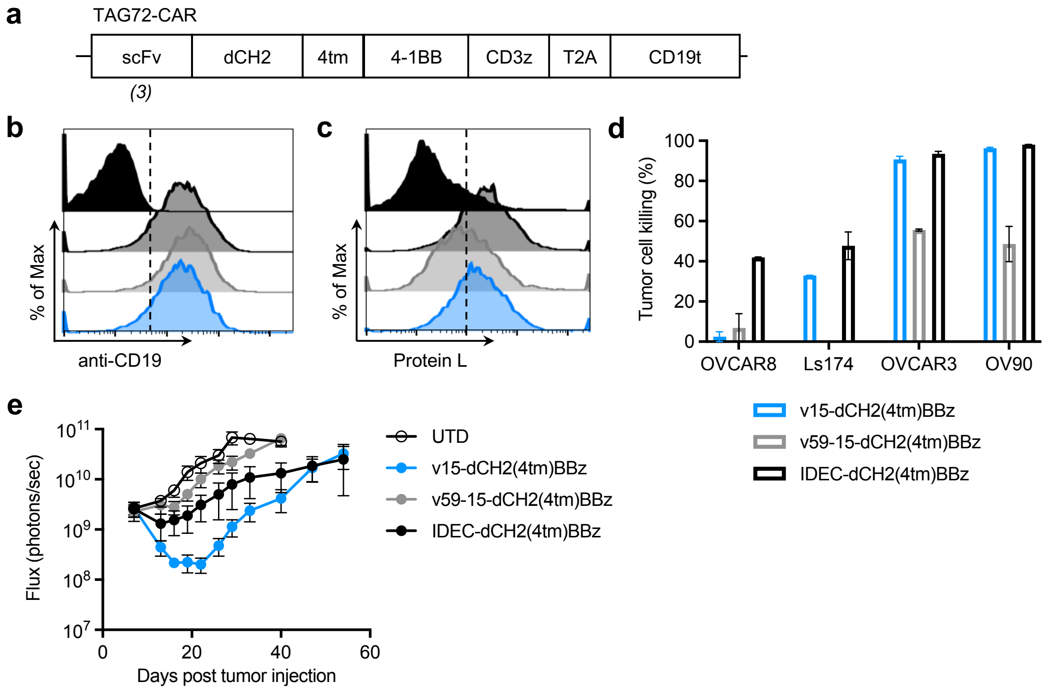


**Supplemental Figure 2. *In vitro* and *in vivo* analysis of TAG72-CAR T cells with varying scFv. (a)** Diagram of the lentiviral construct with a TAG72-CAR containing three different humanized scFvs (based on CC49 clone) targeting TAG72, with dCH2 extracellular spacer domain (dCH2), CD4 transmembrane (4tm), and intracellular 4-1BB costimulatory domain (BBz) followed by a cytolytic domain (CD3z). A truncated non-signaling CD19 (CD19t), separated from the CAR sequence by a ribosomal skip sequence (T2A), was expressed for identifying lentivirally transduced T cells. **(b-c)** Untransduced (UTD) and three TAG72-CAR T cells positively enriched for CD19t were evaluated by flow cytometry for CD19t expression to detect lentiviral transduction of CARs (b), or Protein L to detect the scFv (c). **(d)** Quantification of tumor cell killing by three TAG72-CAR T cells relative to UTD T cells at an E:T ratio of 1:2, following a 72 hour co-culture with antigen-positive and -negative tumor targets as described in Materials and Methods. **(e)** Quantification of flux from i.p. OV90(eGFP/ffluc) tumor-bearing mice treated i.p. with UTD or TAG72-CAR T cells. n = 6-7 per group.

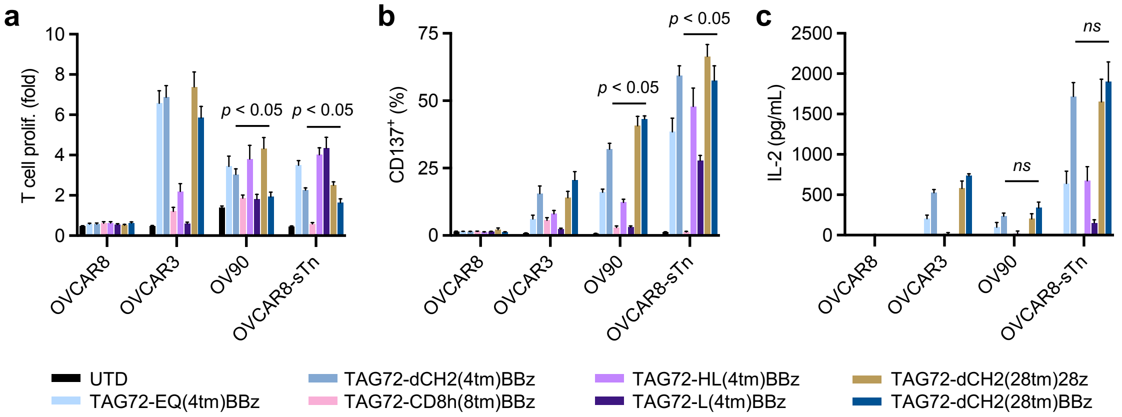


**Supplemental Figure 3 In vitro and in vivo analysis of TAG72-CAR T cells with varying extracellular spacer, transmembrane and costimulatory domains.**

*In vitro* T cell proliferation in fold change compared to UTD (a), expression of CD137 (b), and IL-2 production by ELISA (c), of CAR T cells against tumor targets (TAG72- OVCAR8; TAG72+ OVCAR3, OV90, and OVCAR8-sTn) after 24 hr (for ELISA) or 72 hr of coculture at an effector:target (E:T) ratio of 1:4.

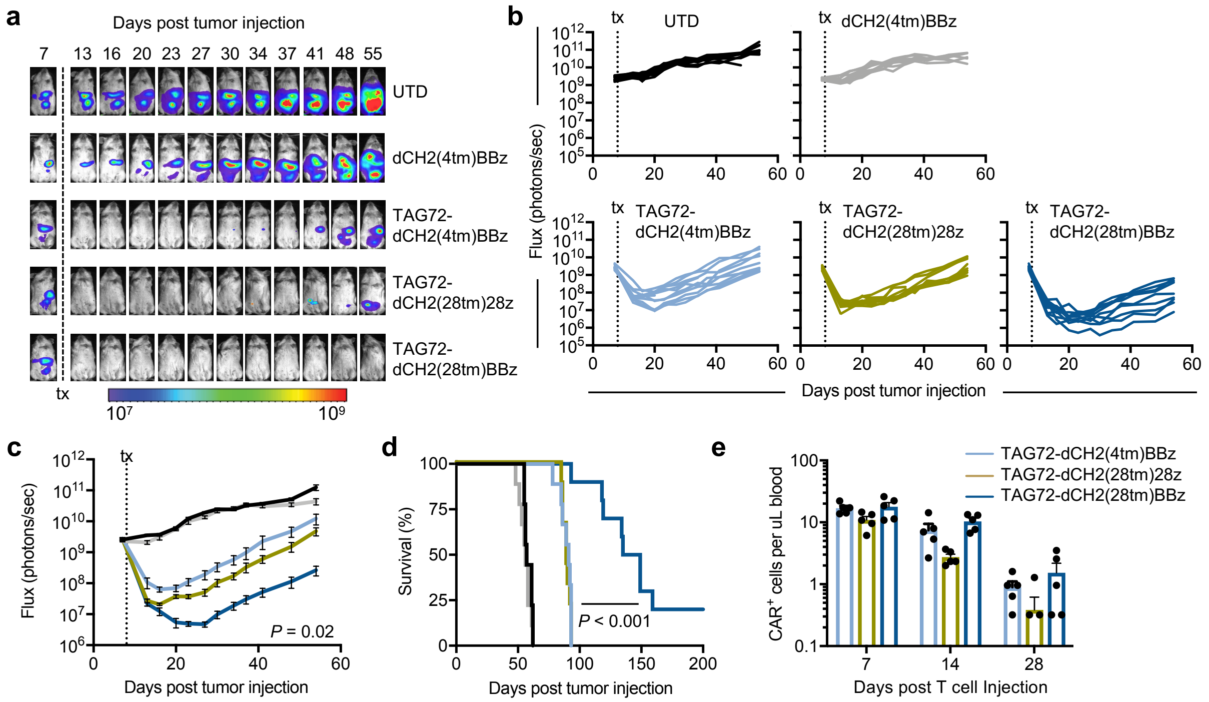


**Supplemental Figure 4. Regional intraperitoneal delivery of TAG72-dCH2(28tm)BBz CAR T cells reduces tumor burden and extends overall survival *in vivo*.**

**(a)** Representative bioluminescent flux imaging of mice treated i.p. with UTD or TAG72-CAR T cells. **(b-c)** Quantification of flux (b, individual mice per group; c, averages) from i.p. OV90(eGFP/ffluc) tumor-bearing mice treated i.p. with UTD or TAG72-CAR T cells. n = 9 mice per group. **(d)** Kaplan-Meier survival for UTD, scFv-less, and TAG72-CAR T cell treated mice. n = 8 - 9 mice/group. **(e)** Quantification of TAG72-CAR T cells per uL of peripheral blood at 7, 14, and 28 days post-treatment. n = 5/group.

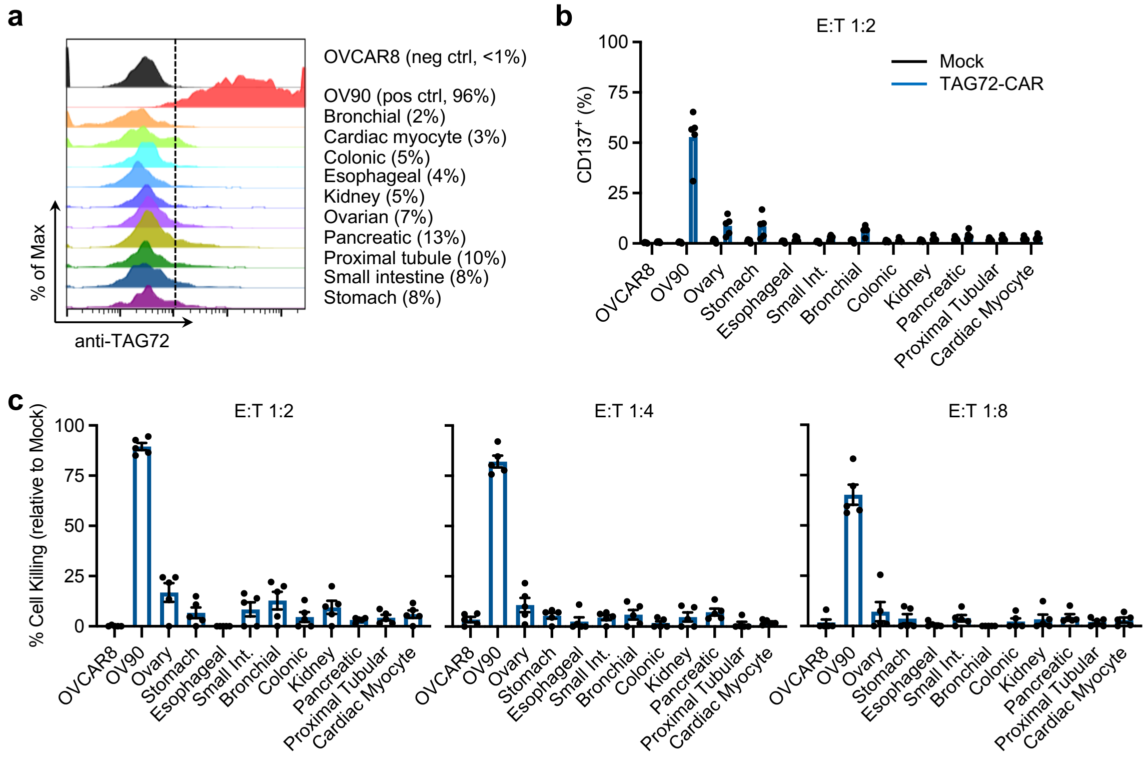


**Supplemental Figure 5. *In vitro* safety of TAG72-CAR T cells against normal human cell lines. (a)** Flow cytometric analysis of TAG72 expression on the cell surface of TAG72-negative (OVCAR8) tumor cells, TAG72-positive (OV90) tumor cells, and indicated primary human normal cells. **(b-c)** Quantification of CD137 activation (b), and tumor and normal cell killing by TAG72-dCH2(28tm)BBz CAR T cells relative to UTD T cells at varying E:T ratios (c), assessed by flow cytometry following a 48 hour coculture with indicated cells as described in Materials and Methods. Data are representative of five independent donors.

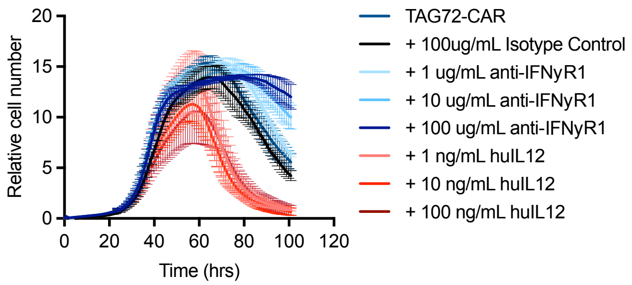


**Supplemental Figure 6. TAG72-CAR T cell anti-tumor activity is regulated by IFNγ signaling.** Tumor cell killing of OVCAR3 cells by TAG72-CAR T cells (E:T = 1:50) with varying concentrations of anti-IFNγR1 blocking antibody, isotype control, and recombinant human IL-12 cytokine measured by xCELLigence.

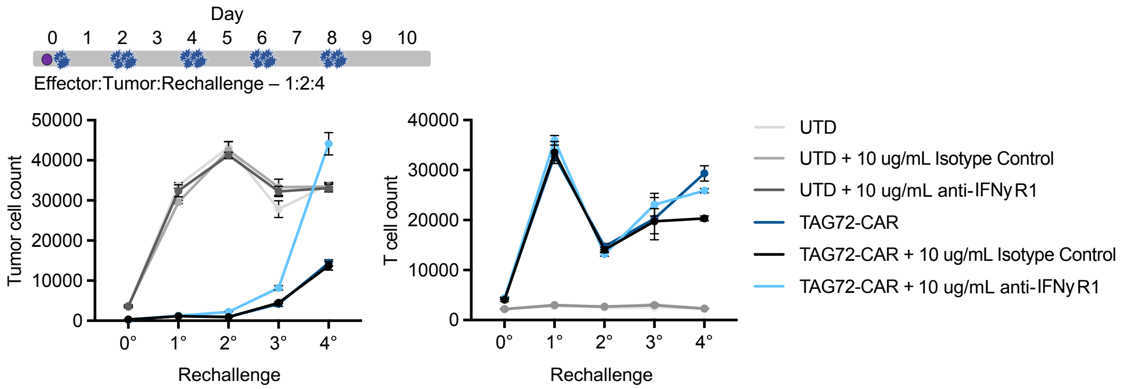


**Supplemental Figure 7. Inhibition of IFNγ signaling dampens repetitive tumor cell killing by TAG72-CAR T cells.** Schema of repetitive tumor cell challenge assay (top). TAG72-CAR T cells were cocultured with OV90 cells (E:T = 1:2) and rechallenged with OV90 cells every three days. Remaining viable tumor cells and TAG72-CAR T cells were quantified as described in Materials and Methods prior to each tumor cell rechallenge.

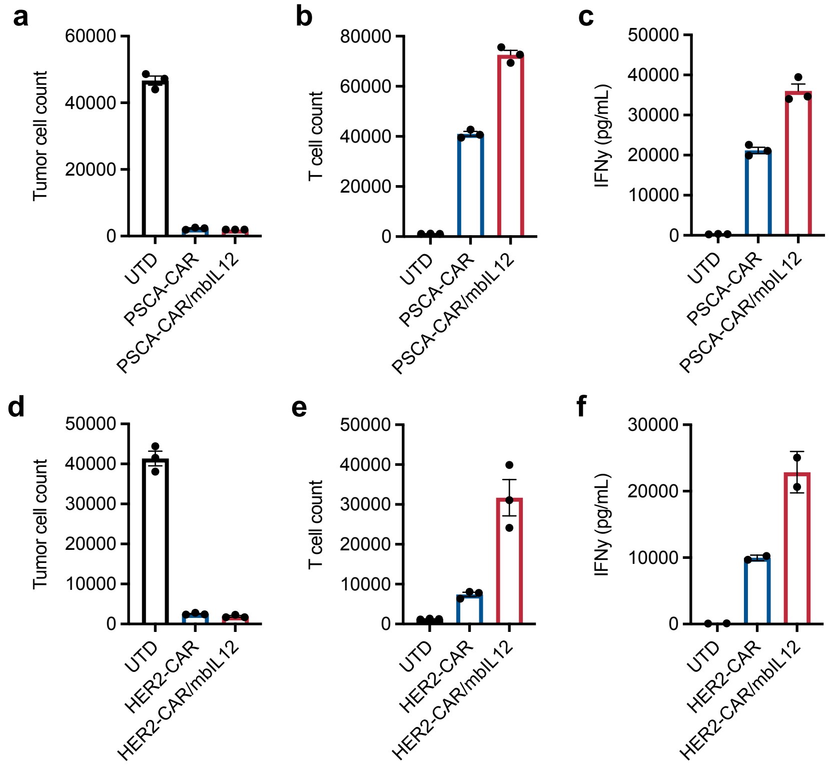


**Supplemental Figure 8. mbIL-12 signaling improves *in vitro* functionality of PSCA- and HER2-CAR T cells. (a-f)** Quantification of tumor cells (left), T cells (middle), and IFNγ levels in supernatant (right) for PSCA-CAR T cells (a-c) and HER2-CAR T cells (d-f), following coculture with antigen-positive targets at an E:T ratio of 1:10 for 6 days.

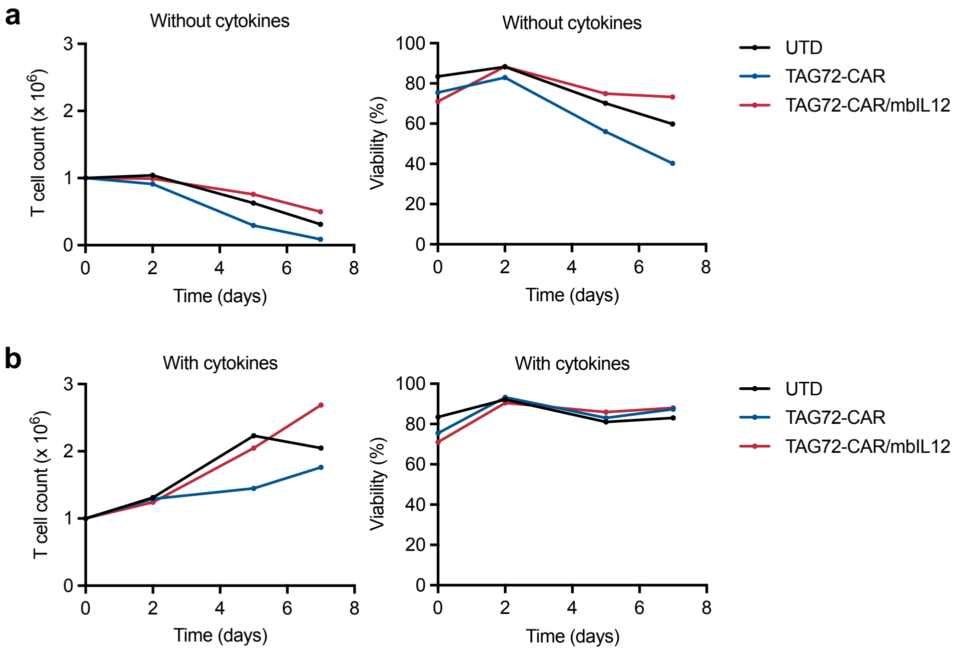


**Supplemental Figure 9. mbIL-12 expressing TAG72-CAR T cells do not expand and survive in the absence of exogenous cytokines. (a-b)** Quantification of T cell count (left) and percentage of viable cells (right) during *ex vivo* culture in the absence (a) or presence (b) of exogenous cytokines as described in Materials and Methods.

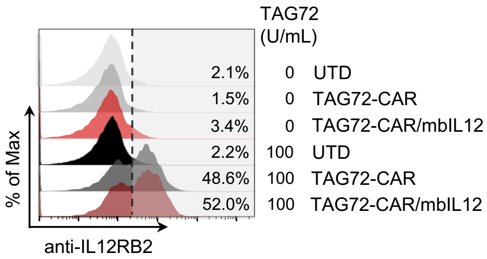


**Supplemental Figure 10. IL12RB2 surface expression is antigen-dependent through CAR-stimulation.**

Flow cytometric analysis of surface expression of IL12RB2 on TAG72-CAR T cells unstimulated or stimulated with plate-bound TAG72.
